# New plasmon-polariton model of the saltatory conduction

**DOI:** 10.1101/2020.01.17.910596

**Authors:** W. A. Jacak

## Abstract

We propose a new model of the saltatory conduction in myelinated axons. This conduction of the action potential in myelinated axons does not agree with the conventional cable theory, though the latter has satisfactorily explained the electrosignaling in dendrites and in unmyelinated axons. By the development of the wave-type concept of ionic plasmon-polariton kinetics in axon cytosol we have achieved an agreement of the model with observed properties of the saltatory conduction. Some resulting consequences of the different electricity model in the white and the gray matter for nervous system organization have been also outlined.

**SIGNIFICANCE:** Most of axons in peripheral nervous system and in white matter of brain and spinal cord are myelinated with the action potential kinetics speed two orders greater than in dendrites and in unmyelinated axons. A decrease of the saltatory conduction velocity by only 10% ceases body functioning. Conventional cable theory, useful for dendrites and unmyelinated axon, does not explain the saltatory conduction (discrepancy between the speed assessed and the observed one is of one order of the magnitude). We propose a new nonlocal collective mechanism of ion density oscillations synchronized in the chain of myelinated segments of plasmon-polariton type, which is consistent with observations. This model explains the role of the myelin in other way than was previously thought.

## INTRODUCTION

Action potential (AP) signal kinetics in dendrites or unnmyelinated axons is well described by the cable theory (1–3). Upon this theory the velocity of an AP signal is characterized by 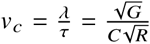, where *C* is the capacity across the neuron cell membrane per unit of the axon length, *G* is the conductance across the membrane and *R* is the longitudinal resistance of the inner cytosol, both per unit of the neuron length, and *λ* and *τ* are space and time diffusion ranges defined in the cable model, respectively (1–5). The velocity *v_c_* scales as 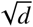 with the dendrite (axon) diameter, *d*, because 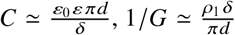 and 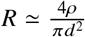, where *δ* is the cell membrane thickness, *ρ* is the longitudinal resistivity of the inner cytosol and *ρ*_1_ is the resistivity across the membrane. For exemplary values of the membrane capacity per surface unit, *c_m_* = 1*μ*F/cm^2^, *ρ*_1_*δ* = 20000 **Ω**cm^2^, *ρ* = 100 **Ω**cm and the diameter of the cable *d* = 2 *μ*m, one gets *v_c_* ≃ 5 cm/s (1).

Due to a myelin sheath in myelinated axons both the capacity and conductance across the myelinated membrane are reduced, roughly inversely proportional to the myelin layer thickness. Thus, the cable theory velocity, 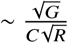, grows ca. 10 times if the capacitance and conductance lower ca. 100 times. This is, however, still too low to match observations of kinetics in myelinated axons (1–4, 6–12). Moreover, the activity of nodes of Ranvier slows down the overall velocity of the discrete diffusion in the myelinated internodal segments to the level of factor 6 instead of 10 (1). For realistic axons with *d* ~ 1 *μ*m the assessed cable theory velocity gives ca. 3 m/s instead of 100 m/s observed in such-size myelinated axons in peripheral human nervous system (PNS). Similar inconsistency in the cable model estimations is encountered in human central nervous system (CNS) with thinner myelinated axons of *d* ∈ (0.2,1) *μ*m (6).

To model larger velocities upon the cable theory accommodated to the myelinated axons, much thicker axons are assumed with longer internodal distances, because upon the discrete diffusion model (1), the velocity in a myelinated axon, 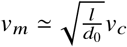, where *l* is the length of internodal segments and *d*_0_ is the length of the node of Ranvier. For instance, in (6, 8) *d* = 10*μ*m (and greater) and *l* = 1150 *μ*m (and greater) have been assumed to gain the velocity of the AP transduction, *v_m_* ~ 40 m/s (cf. also (9–12)), i.e., there are ca. ten times larger in dimension than the actual myelineted axons of a human. It is thus clear that the cable model is not effective in modeling of the observed quick saltatory conduction in myelinated axons (2–4, 6–13). The velocity predicted upon the discrete cable model for realistic axon parameters is by at least one order of the magnitude smaller than observed.

Interfering of the Huxley-Hodgkin (HH) mechanism at nodes of Ranvier with the cable model diffusion at internodal myelinated segments results in the estimation of the AP propagation velocity (9, 11) only ca. 6 time greater than in unmyelinated axons with the same geometry (1–3), despite the reducing of the intercellular capacity and conductivity by the myelin sheath. The mentioned above simplified formula for this velocity (for iterative discrete diffusion model (1)) gives 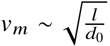, and since *d*_0_ is often 1 *μ*m and *l* is around 100 *μ*m, the increase in velocity of myelinated axons can be almost 10 times that of unmyelinated axons (more precisely the factor is closer to 6 (1)).

Velocity of the AP transduction must keep the high value because the deviation by 10% ceases body functioning (6). Continued mathematical attempts to optimize a model of HH mechanism mixed with the cable theory (1–3, 9, 11) in order to obtain a sufficiently high AP velocity in myelinated axons gave not a success for a long time, which strongly suggest that the way to understand the saltatory conduction must be linked with a different physical mechanism rather than any lifting of the cable theory of ion diffusion.

In this paper we propose a quite new approach to the problem of the saltatory conduction in myelinated axons by a wave type of signaling of the ion plasmon-polariton with physical mechanism and properties different than that of the ordinary charge diffusion current of ions. We have achieved the compliance with the most important and puzzling properties of the saltatory conduction not attainable for the cable theory. Below we present a relevant mathematical model which allows for the quantitative verification of the plasmon-polariton ion kinetics against real myelinated axon parameters and saltatory conduction features including those inaccessible for the conventional ion-diffusion current picture but possible to be understood upon a concept of the collective wave-type synchronized oscillations of ion density in the periodic chain of myelinated segments.

## PLASMON-POLARITON WAVE-TYPE CONCEPT FOR THE SALTATORY CONDUCTION

We propose to abandon the cable theory and assume a different scenario for signal transduction along the myelinated axons. The HH mechanism at nodes of Ranvier coupled with a discrete diffusion along adjacent internodal myelinated segments constitutes the local electrical approach. We propose an essentially nonlocal mechanism in the form of the ionic plasmon-polariton wave-type collective propagation along the whole axon periodically corrugated by the myelin sheath. Instead of a local longitudinal diffusion current of ions in the axon cord, we propose the synchronized oscillations of ion density in consecutive internodal segments propagating in the wave manner along the whole axon. The wave packet of such synchronized oscillations, i.e., of the ionic plasmon-polariton in analogy to plasmon-polaritons in metallic linear nano-structures (14–18), may serve as a carrier of the ignition potential beyond the voltage threshold required for the triggering of the HH cycle on consecutive nodes of Ranvier to fire the whole axon. This transmission of the ignition voltage undergoes without any net diffusion current of ions and its velocity is the group velocity of the plasmon-polariton wave packet not limited by the large longitudinal resistivity of the inner cytosol of the axon cord. The wave type propagation of synchronized oscillations of ion density in all segments of the myelinated axon is allowed due to periodicity imposed by the dielectric myelin sheath and the propagation of the wave packet of plasmon polariton undergoes with the velocity conditioned by the frequency of ion plasma oscillations in iternodal myelinated segments and by the geometry, size and physical parameters of the axon and of the myelin sheath. This velocity is not reduced by a poor longitudinal conductivity of inner cytosol of axons. The ion density oscillation frequency is also itself dependent on the geometry-size, and on material parameters of the axon cord and of the myelin sheath. The thickness of the myelin cover and its periodicity is of the primary importance. The insulating dielectric myelin sheath influences the AP kinetics in a different way in comparison to the cable theory model.

If the amplitude of oscillating dipoles synchronized along the myelinated segments exceed the threshold needed to initiate HH cycle at a particular node of Ranvier, then this node begins its HH activity and the local voltage of the AP spike attains ca. six times larger value due to an influx of Na^+^ions through open trans-membrane channels in time period less than 1 ms. This local increase of the polarization at the active node of Ranvier supports residually the mode of plasmon-polariton with such a frequency which is synchronized with the timing of the Na^+^channel gating. Too large frequency of ion density oscillations precludes opening of the gate (the timing of the gate opening is of the scale of microseconds) but too small frequency inconveniently lowers the velocity of the ionic plasmon-polariton, on the other hand. Thus, via the trade-off of the opposite factors, the HH cycle plays the regulatory role imposed on the plasmon-polariton and supplements in energy the synchronized mode of the plasmon-polariton wave-packet. Unsynchronized modes, not supplemented in energy, are quickly quenched on short distances because of the Ohmic energy losses due to ion scattering at oscillations on other ions, on admixtures and on solvent molecules. We have verified that the plasmon-polariton mode synchronized with the HH cycle propagates with the group velocity of 100 – 200 m/s for the wave packet which carries a voltage signal effectively activating consecutive nodes of Ranvier. Moreover, the plasmon-polariton wave packet synchronized with the HH cycle is conveniently step-by-step narrowed by HH cycles at consecutive nodes of Ranvier via the distributed in time-space continuous additional energy supporting, but only of synchronized plasmon-polariton mode (each HH cycle adds energy and narrows the wave packet of the plasmon polariton).

In a time period of 1 ms, the plasmon-polariton wave packet initiates ca. 1000 nodes of Ranvier along 10 cm of the myelinated axon (with *l* = 100 *μ*m length of the myelinated segment and for plasmon-polariton packed speed, *v_m_* = 100 m/s). Within this 1 ms the ignited one-by-one consecutive nodes of Ranvier are on various steps of their individual HH cycles, as the whole HH cycle takes ca. 1 ms (and longer, even up to a second is needed for restoring of the steady state of Ranvier node, which disactivates temporarily a node of Ranvier and blocks reversing of the plasmon-polariton wave). Plasmon-polariton is not a local effect, it propagates along the whole myelinated axon and is conditioned by the myelination periodicity scale (given by *l*—the length of a myelinated segment). The length of the myelinated internodal segments, the myelin thickness and the width of Ranvier nodes play a regulatory role similarly as the diameter of the axon rod. In the next paragraph we define the mathematical model of the ion plasmon-polariton kinetics along a myelinated axon and verify their characteristics versus the real geometry and physical material parameters of the axon cytosol and the myelin cover.

Plasmon-polariton (14–18) is a hybridization of surface plasmon oscillations (19–21) with electromagnetic (e-m) field, thus it can propagate despite small discontinuity of the axon (22). This exceptional property explains the observed maintenance of myelinated axon firing even if the axon is broken into two parts with the ends separated slightly (4). Such a gap is impossible to be overcome by the diffusion current, but the small gap is not an obstacle for the plasmon-polariton (14–18). Similarly, a damage of several nodes of Ranvier (either grouped or randomly distributed) only weakly disturbs the plasmon-polariton kinetics but completely prohibits the discrete diffusion ion current upon the cable model. The group velocity of the plasmon-polariton depends significantly on the thickness of the myelin sheath. Any deficiencies in the myelin cover cause the reduction of plasmon-polariton speed. The myelin sheath plays here a double role. Besides imposing the periodicity, it also creates the dielectric tunnel between the inner and outer cytosol electrolytes unavoidably required for the plasmon-polariton formation. The size and geometry of this channel is a crucial factor. These facts together with the consistency of the plasmon-polariton behavior with the observed saltatory conduction in myelinated axons (including a temperature and size dependence) favor strongly the proposed model. Moreover, the AP transduction in the scenario of the plasmon-polariton kinetics is very energy frugal (more than for the diffusion current) and reliable over arbitrary long distances. Plasmon-polariton does not emit any e-m radiation energy (14–18)—all the e-m field is compressed in its own structure along the axon cord. The plasmon-polariton is radiatively lossless in contrary to the ordinary diffusion current. Plasmon-polariton cannot be neither detected nor perturbed by an external e-m field—thus, the saltatory conduction in myelinated axons is especially robust against all external e-m perturbations, which enhances the reliability of this information channel for signaling in both PNS and CNS, most important for the body functioning.

## MATHEMATICAL DESCRIPTION OF PLASMON-POLARITON KINETICS IN THE CHAIN OF MYELINATED SEGMENTS OF AN AXON

We propose to apply the model of dipole plasmon-polariton excitations in a linear chain of electrolyte segments to an axon cord periodically wrapped with the myelin sheath, as schematically depicted in Fig. 1. The periodicity of these segments makes such a system similar to a 1D crystal. Despite the cord of an axon is a thin continuous tube filled with the electrolyte, the modeling of an axon by a chain of segments defined by the periodic myelin sheath meets well with plasmon-polariton kinetics maintaining the same character in discrete chains and in continuous but periodically corrugated conductor wires—the property very well known in plasmonics (23–26). The interaction between the chain segments (or segments defined by the myelin sheath in the axon cord) can be regarded as the dipole coupling of local ion oscillations in each segment, similarly as for electron oscillations in metallic nano-chains (14–16, 27, 28). Ions have larger mass than electrons (ca. 10^4^ times) and a concentration in a cytosol much lower than of electrons in metals, hence their plasmon oscillation frequencies (thus energies) are far lower that those of metallic electron plasmons. It changes also the so-called plasmonic size (defined as the most convenient for plasmon-polariton formation) from nanometer scale for electrons to micrometer scale for ions. The size and energy scale of plasmon-polariton in ionic systems matches with that required in bio-living system, just as in myelinated axons.

**Figure 1:**
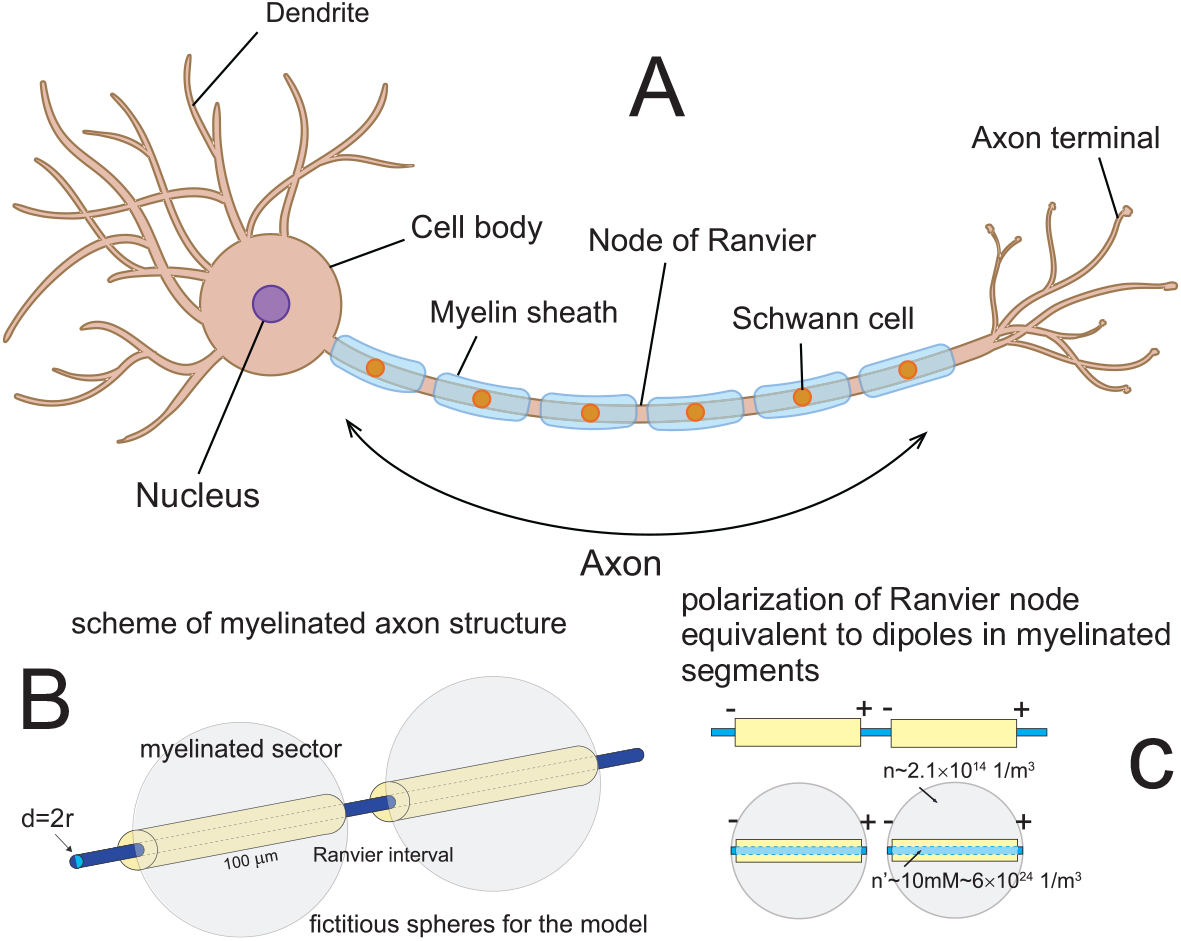
In analogy to metallic nano-chains, micro-chains of finite ionic segments confined by dielectric membranes and periodically myelinated (A) can be considered; in these ionic micro-chains the plasmon-polaritons can traverse both continuous and discrete linear plasmonic system alignments; polarization of the node of Ranvier is given by oscillating dipoles of the plasmon poriton propagating along the chain of myelinated segments (C); fictitious auxiliary spheres used for the estimation of longitudinal surface plasmon oscillation frequency in myelinated segments (B,C).

The dipole interaction between segments with oscillating ions resolves itself to the electric and magnetic fields created by the oscillating dipoles at each segment of the chain. The oscillating dipole, **D**(**r**, *t*), with its center in the point **r** (identified as the center of a myelinated segment) induces at any distant point a varying electric and magnetic fields. If this distant point is represented by the vector **r**_0_ (beginning ar **r**, where the dipole is fixed), then the electric and magnetic fields produced by the dipole **D**(**r**, *t*) take the following form, including the relativistic retardation (due to the finite velocity of e-m signal propagation) (29, 30),

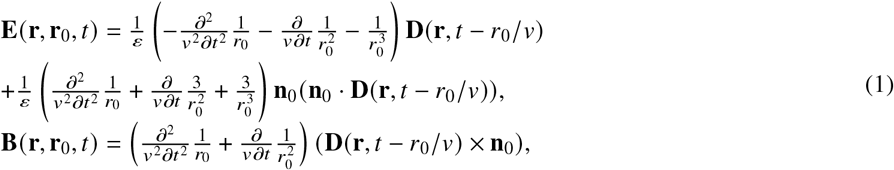

with 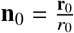 and 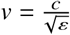 (*ε* is the dielectric permittivity of the medium, the magnetic permeability, *μ* = 1 here, *c* is the light velocity in the vacuum). The terms with denominators of 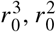, and *r*_0_ are usually referred to as the near-field, medium-field, and far-field zones of the dipole interaction, respectively (29, 30). The above formulae describe the mutual interactions of the plasmon dipoles synchronically distributed over segments in the chain. The electric field dominates strongly (29, 30) this interaction, thus the magnetic field can be neglected here.

The segments in the chain can be numbered by integers *n*. In the chain geometry the origin of the reference coordinate system can be located at a selected chain component—let say, at *n*-th one, then for the dipole located on this segment the vector **r** = 0 and vectors **r**_0_ to other chain segments are collinear and have lengths, *r*_0_ = |(*m* − *n*)*h*|, *m* = ±1, ±2, ±3,… (*h* is separation between neighboring segment centers)—cf. Fig. 2.

**Figure 2:**
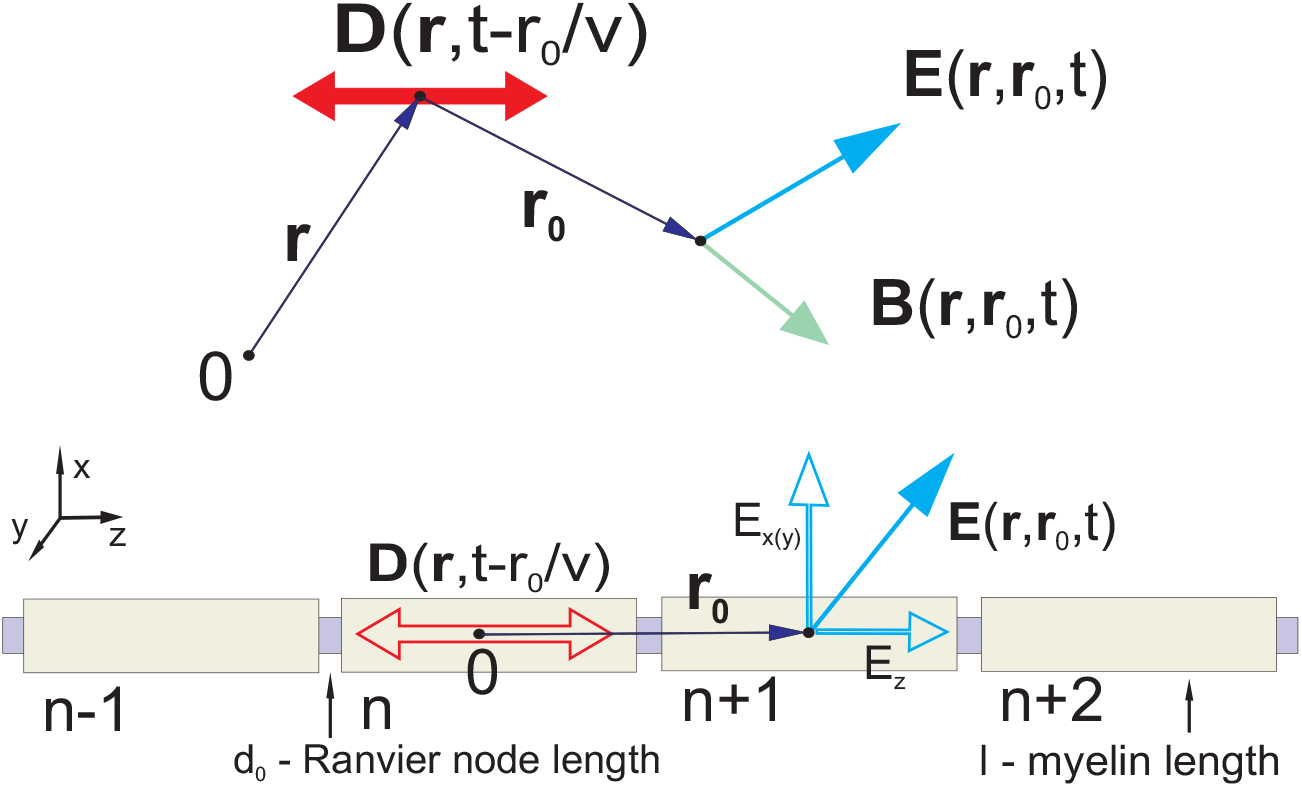
(Upper) Illustration to Eqs (1), the dipole oscillating at point **r** induces electric and magnetic fields at any distant point **r**_0_, with the retardation effect related to a finite velocity of e-m signals, 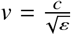. (Bottom) In the axon geometry, four consecutive segments are shown with the origin of the coordinates (0) at *n*-th segment. Vectors to centers of other chain segments, **r**_0_ = (*n − m*)*h ẑ*, are collinear (*ẑ* is the versor of the *z* axis, *h* is the separation between neighboring segment centers, *m* is the number of a segment, *m* ≠ *n*). The length of the node of Ranvier, *d*_0_, is much shorter than the myelinated segment length, *l*, thus the separation between the centers of the chain segments, *h* = *l* + *d*_0_ ≃ *l*. In centers of segments there are placed dipoles of local ion oscillations. These dipoles are coupled according to Eq. (4).

The equation for the surface plasmon oscillation of the particular *n*-th segment can be written as follows (the separation between the centers of neighboring segments *h* ≃ *l*, as in our case the myelinated segments of length *l* are separated by small nodes of Ranvier and *h* = *l* + *d*_0_ ≃ *l*, because *d*_0_ ≪ *l*, cf. Fig. 2),

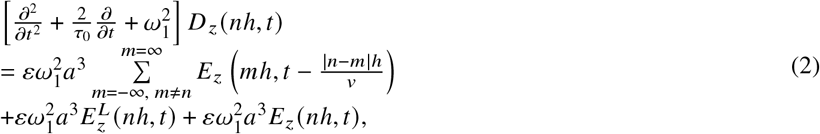

*z* indicates the longitudinal polarization (the chain orientation is assumed to be along the *z* direction), *a* = *l*/2 is the radius of the sphere with diameter *l*. The first term in the right-hand side of Eq. (2) describes the dipole coupling between the *n*-th segment with other segments in the chain (denoted by *m, m* ≠ *n*), and the other two terms correspond to the plasmon attenuation due to the radiation losses, so-called Lorentz friction (29, 30) and the force field arising from an external electric field, respectively; *ω*_1_ is the frequency of the longitudinal dipole surface plasmon in the segment, when it is separated. In a separated segment the Lorentz friction is very large at plasmonic its size (31, 32), but in a chain of segments this large energy losses are balanced by the energy income form other segments if they oscillate in a synchronized manner (18). Besides the radiation one must include also the Ohmic losses due to scattering of oscillating ions with other ions, admixtures, solvent particles in an electrolyte and on walls of the segment. Ohmic losses, eventually transformed into Joule heat, are included here via the term, 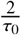, similar to what is applied to metals (31) but with the Fermi velocity of electrons in metals substituted by the mean velocity of ions for classical Boltzmann distribution regardless of the quantum statistics of ions (19), i.e.,

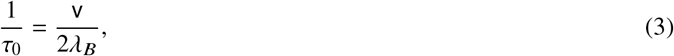

where *λ_B_* is the mean free path of ions the same as in the bulk electrolyte, v is the mean velocity of ions at temperature *T*, 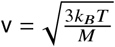, *M* is the mass of the ion, *k_B_* is the Boltzmann constant. The expression for 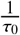 approximates ion scattering losses such as those occurring in the bulk electrolyte (collisions with other ions, solvent and admixture molecules, for longitudinal ion oscillations in a segment of the axon cord, the scattering of ions on the system boundary does not contribute due to axon cord continuity).

According to Eq. (1), we can write out the following quantities that appear in Eq. (2) (for the longitudinal polarization mode, i.e., for *z* polarized ion oscillations),

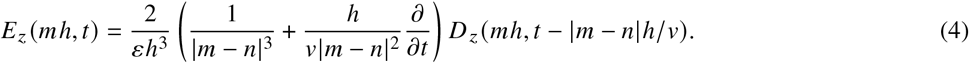

The Lorentz friction term has the form, 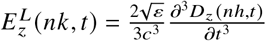 (19, 29, 30, 32). The external forcing field *E_z_* describes the exciting of plasmon-polariton, and as usual for linear differential equations the nonhomogeneous term (like with *E_z_* in Eq. (2), which is the linear equation with respect to *D_z_*) can be avoided if we look for resonance self-frequencies, but is important when a wave packet is shaping.

Because of the periodicity of the chain of myelinated segments, a wave-type collective solution of the dynamical equation (2) in the form of Fourier component can be assumed (as usual for linear differential equations),

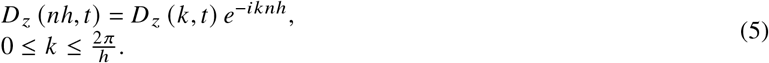

In the Fourier picture of Eq. (2) (the discrete Fourier transform (DFT) with respect to the positions and the continuous Fourier transform (CFT) with respect to time) this solution takes a form similar to that of the solution for phonons in 1D crystals. Note that DFT is defined for a finite set of numbers; therefore, we consider a chain with 2*N* + 1 elements, i.e., a chain of finite length *L* = 2*Nh*. Then, for any discrete characteristic *f*(*n*), *n* = −*N*,…, 0,…, *N* of the chain, such as a selected polarization of the dipole distribution, we must consider the DFT picture 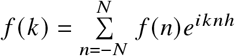, where 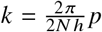, *p* = 0, …, 2*N*. This means that *kh* ∈ [0, 2*π*) because of the periodicity of the equidistant chain. The Born-Karman periodic boundary condition, *f*(*n* + 2*N*) = *f*(*n*), is imposed on the entire system, resulting in the form of *k* given above. For a chain of infinite length, we can take the limit *N* → ∞, which causes the variable *k* to become quasi-continuous, although *kh* ∈ [0, 2*π*) still holds.

The Fourier representation of Eq. (2) takes the following form,

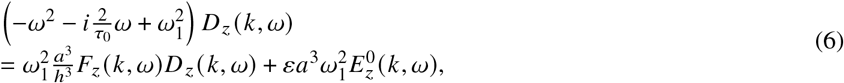

with

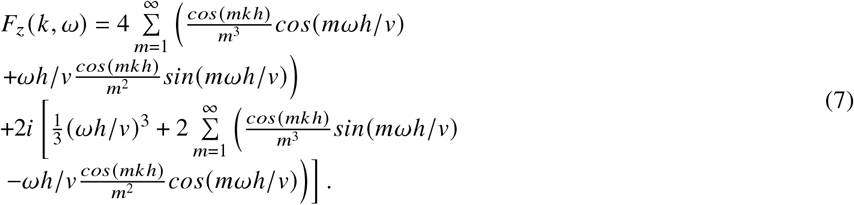

The most important and surprising feature is that *ImF_z_*(*k*, *ω*) ≡ 0, which indicates the perfect quenching of the radiation losses at any segment in the chain. This means that to each segment, the amount of energy that comes from the other segments is the same as the energy outflow due to the Lorentz friction from this segment. We can easily verify this property directly, as the related infinite sums in the imaginary part of Eq. (7) can be found analytically (33). These sums have the following form,

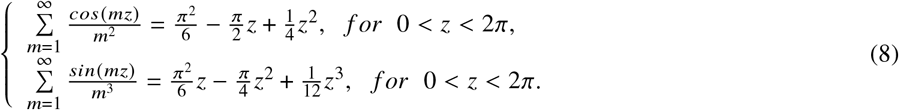

To be more specific, let us note that,

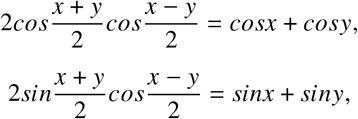

which allows for an analytical calculation only of the imaginary part of Eq. (7) (with 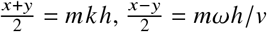) by virtue of Eqs (8) with *x, y* substituting *z* appropriately (because Eqs (8) hold for 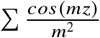 and 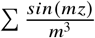, but not for 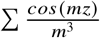 and 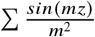, two latter sums are not given analytically).

Eq. (6) is highly nonlinear with respect to the complex *ω* and can be solved both perturbatively in an analytical manner (17) or numerically, which is more accurate (18). The solutions determined for *Reω* and *Imω* give the self-frequency of the plasmon polariton and its damping, respectively. The *k*-derivative of the real part of the resonance frequency (*Reω*(*k*)) defines the group velocity of particular mode *k*. The imaginary part of the resonance frequency (*Imω*(*k*)) defines the damping of the dipole plasmon-polariton *k* mode. The verified above property, that *ImF_z_* (*k, ω*) ≡ 0, means that the plasmon polariton does not lose e-m energy and its damping is caused only by relatively small Ohmic losses, hence this kinetics is energy frugal. This surprising property indicates that all large energy radiated by plasmons (due to Lorentz friction) is perfectly balanced by energy income from other segments in the chain at synchronized propagation along the chain of wave-type ion oscillations, and hence none energy is emitted radiatively from the chain by plasmon-polariton.

The problem of propagation of plasmon-polaritons along metallic nano-chains was developed, among others, by Citrin (34, 35) in order to assess the radiative losses of plasmon-polaritons in metallic nano-chains in agreement with former observations related to radiation losses in one-dimensional and two-dimensional crystals (36). For more detailed calculations, including also the transverse polarization of plasmon-polariton modes and the formation of a plasmon-polariton wave packet, cf. (17, 18, 22, 37).

## FITTING THE PLASMON-POLARITON KINETICS TO THE AXON PARAMETERS

The bulk plasmon frequency for ions in an electrolyte (19, 20) is 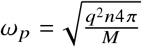 (we assumed for the model that the ion charge is *q* = 1.6 × 10^-19^ C and the ion mass is *M* = 10^4^*m_e_*, where *m_e_* = 9.1 × 10^-31^ kg is the mass of an electron) and for a concentration of *n* = 2.1 × 10^14^ 1/m^3^, we obtain the Mie-type frequency for ionic dipole oscillations, 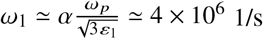, where the relative permittivity of water is *ε*_1_ ≃ 80 for frequencies in the MHz range (38) (though for higher frequencies, beginning at approximately 10 GHz, this value decreases to approximately 1.7, corresponding to the optical refractive index of water, 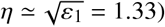). The parameter *α* ≪ 1 describes deviation of the dipole longitudinal oscillation frequency in highly elongate segment of a thin axon cord from the Mie-type frequency value for a sphere, 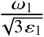. For a strongly prolate inner ionic cord segment, the longitudinal Mie-type frequency is renormalized by a factor of *α* dependent on the aspect ratio (39, 40).

**Figure 3:**
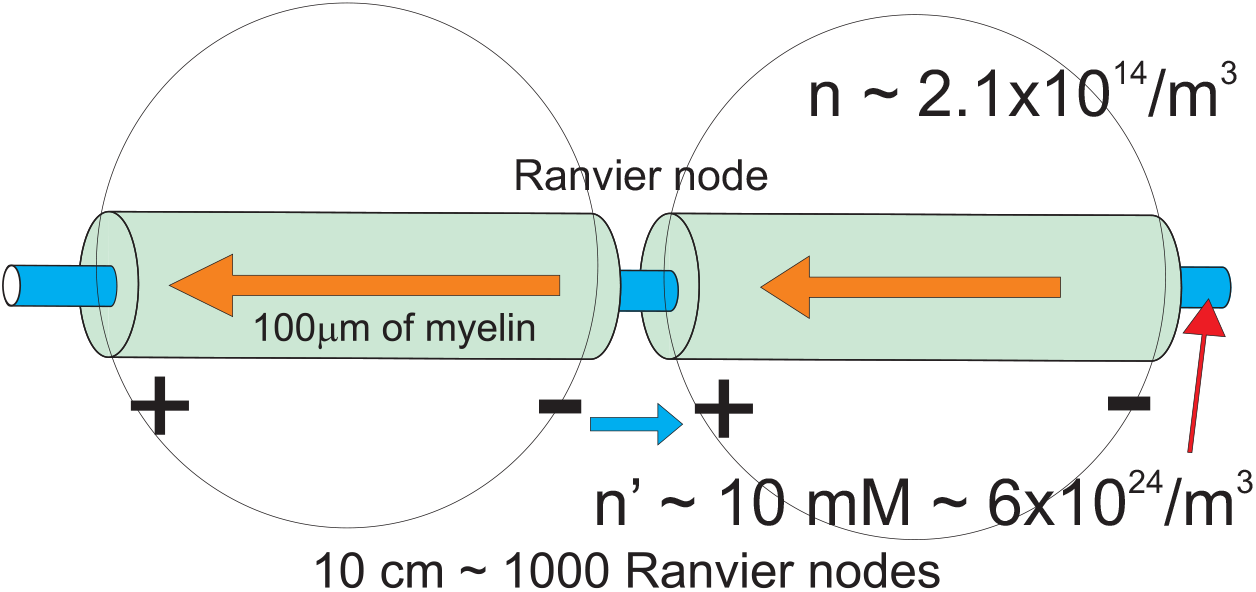
For a model estimation we consider spheres equivalent to periodic myelinated segments with the same number of ions as in the corresponding fragment of the axon rod. The concentration of ions in fictitious spheres for an effective model of finite electrolyte chain of spheres is strongly reduced which gives the required group velocity (ca. 100 m/s) of the ionic plasmon-polariton wave packet. Longitudinal dipole oscillations of surface plasmons in consecutive myelinated segments changing along the axon according the running wave with velocity 100 m/s (red arrows) are equivalent with opposite polarization of Ranvier nodes (blue arrow).

The axon consists of a cord with a small diameter of *d* = 2*r*, and this thin cord is wrapped with a myelin sheath of a length of *l* = 2*a* per segment. For the effective model, one can consider fictitious electrolyte spheres of radius *a*. Thus, the auxiliary concentration n of ions in the fictitious sphere corresponds to the ion concentration in the cord of 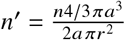, which yields a typical concentration of ions in a nerve cell of *n′* ~ 10 mM (i.e., ~ 6 × 10^24^ 1/m^3^)—cf. Fig. 3. This is because all the ions participating in the dipole oscillation correspond in the sphere model to a much smaller volume in the real system, that of the thin cord segment (the insulating myelin sheath consists of a lipid substance without any ions). The insulating, relatively thick myelin coverage creates the periodically broken channel (corrugated conductor-insulator structure) required for plasmon-polariton formation and its propagation. To reduce the coupling with the surrounding inter-cellular electrolyte and protect against any leakage of plasmon-polaritons, the myelin sheath must be sufficiently thick, much thicker than what is required merely for electrical insulation. The Mie-type frequency of the longitudinal oscillations in the myelinated segment is lower by *α* than that for a sphere with a diameter equal to the elongation axis. As a rough estimate, we have assumed a correction factor of 0.1 (39, 40), which agrees with the presented below assessment of frequency of beating due to coupling across the myelin layer.

To describe the regulatory role of the myelin layer thickness let us consider a single myelinted segment of length *l*. By *d*_1_(*t*) we denote the longitudinal dipole in the axon rod in this myelinated segment. *d*_2_ is the dipole in the outer cytosol nearby adjacent to the segment and induced by *d*_1_ dipole—oppositely directed. *d*_2_ dipole is activated by *d*_1_ and vice verse. Equation for the dynamics of this sub-system is as follows,

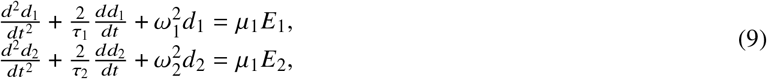

where, 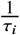 is the damping rate of the surface plasmon *i*-th dipole, *ω_i_* its self-frequency, *μ_i_* is the longitudinal polarizability of the rod for *i* = 1 and of the cave in surrounding cytosol for *i* = 2 (the polarizability for a sphere equals to 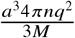, *a*—the sphere radius, *n*—concentration of ions, *M*—ion mass, *q*—ion charge). The electrical fields induced by dipoles in a near-field zone are, 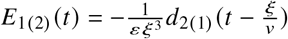, where *ξ* is the thickness of the myelin sheath, 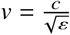, and these fields describe mutual interaction of dipoles *d*_1_ and *d*_2_, i.e., the electrical induction caused by opposite dipoles in the near-field coupling approximation. Simplifying (for illustration) by assuming *μ*_1_ = *μ*_2_ = *μ*, *ω*_1_ = *ω*_2_ = *ω*_0_ and neglecting here damping and retardation, Eq. (9) attains the shape,

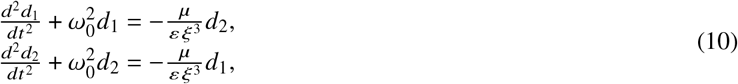

This is the equation for two coupled harmonic oscillators. It has the self-frequencies, for assumed solution, *d*_1_(*t*) = *Ae*^*i*Ω*t*+*ϕ*^ and *d*_2_(*t*) = *Be*^*i*Ω*t*+*ψ*^, given by,

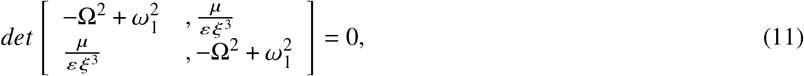

i.e., 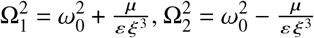.

For the initial condition suitable for excitation of the considered segment of the axon, i.e., *d*_1_(0) = *D*, *d*_2_(0) = 0, 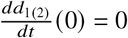, one gets the beating with low frequency, 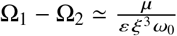. For sufficiently large *ξ* we get thus slow oscillations required for the time scale of opening of the Na^+^ ion channels at nodes of Ranvier to trigger the HH cycle. The period of the igniting signal (with an amplitude beyond the threshold for Na^+^ ion channel opening) cannot be lower than the characteristic time of the activation of these ion channels.

The frequency for longitudinal plasmon is reduced in strongly elongated axon rod segment with the aspect ratio ~ 10^-3^ and additionally reduced by an appropriate increase of *ξ* due to beating effect and finally achieves the value of ~ 10^6^ 1/s, resulting in the saltatory conduction velocity ~ 100 m/s. Thus we see, that the myelin layer thickness controls the velocity of the saltatory conduction and simultaneously accommodates the frequency of the igniting signal oscillation (of plasmon-polariton wave packet) to the time scale of triggering of the ion channels at nodes of Ranvier (being of order of a microsecond). Only this frequency is selected and the corresponding plasmon-polariton wave packet is strengthened by synchronized HH cycles in contrary to other non-synchronized frequency modes of the plasmon-polariton. Nonsynchronized modes are quickly damped due to Ohmic losses.

At the Ranvier node the inner and outer cytosol are separated by the thin bare cell membrane, thus dipole coupling across the thinner barrier is stronger and the corresponding beating quicker. The quick component of this beating is also quenched as not synchronized with the time scale of electrically gated Na^+^ channels, in contrary to slow component of the trans-myelin beating. Via synchronization with HH cycle ignition time scale, this selected mode of plasmon-polariton is continuously supplemented in energy by HH cycles and simultaneously is able to ignite the HH cycle on consecutive Ranviere nodes along the chain of myelinated segments. This explains why only the myelinated segments oscillate and the frequency of related plasmon-polariton wave packet is perfectly tuned by myelin thickness. The oscillation of longitudinal surface plasmon on the myelinated segment of axon rod is equivalent with opposite dipole oscillation (polarization) of the node of Ranvier (but it is not its own self-oscillation). This is schematically illustrated in Figs 4 and 3.

**Figure 4:**
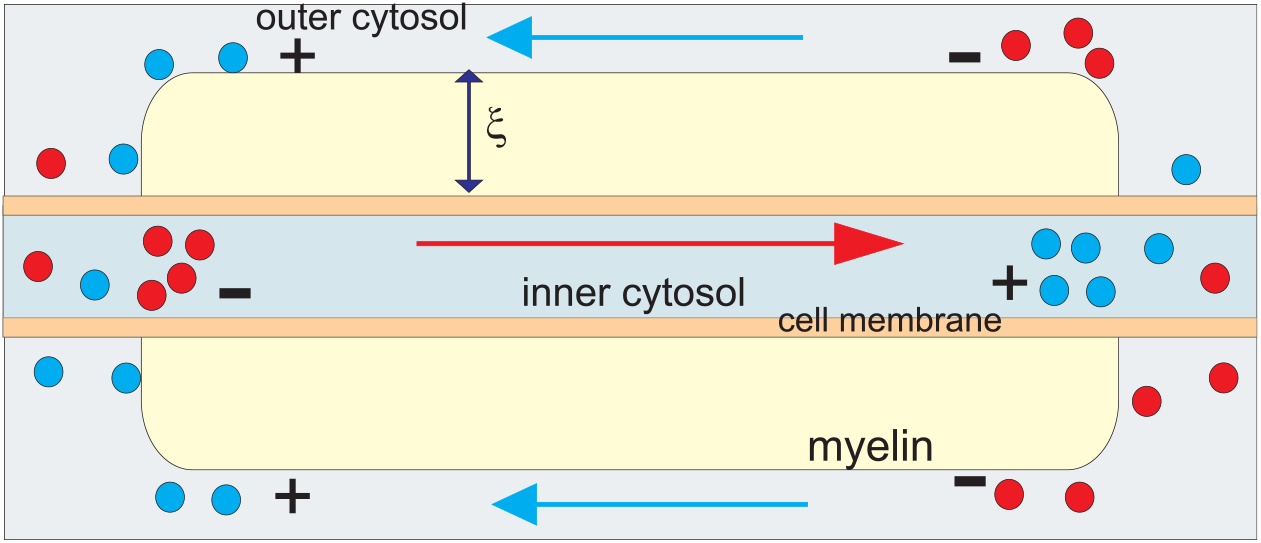
Cartoon of a single myelinated segment—polarization of longitudinal dipole type of the inner cytoplasm inside the axon cell (red arrow) induces local opposite polarization of the outer cytoplasm (blue arrows). Both dipoles oscillate as the pair of coupled oscillators across the insulating myelin layer. The coupling is weak for sufficiently thick myelin sheath and causes slow beats due to oscillator coupling. At the nodes of Ranvier the coupling across much thinner cell membrane is strong and causes quick beating out of a resonance with ion gate timing, thus damped. Slow beating oscillations of coupled dipoles are ranged only to the myelinated sector.

For the resulting frequency of plasmon oscillation of myelinated segments, *ω*_1_ ≃ 4 × 10^6^ 1/s, one can determine the plasmon-polariton self-frequencies in the chain of segments with the length *l* = 2*a* = 100 *μ*m and for small chain separations of *d*_0_/*a* = 2.01, 2.1, and 2.2 (corresponding to node of Ranvier lengths of 0.5, 5, and 10 *μ*m, respectively) within the approach presented above (via the solution of Eq. (6)). The derivative of the obtained self-frequencies (the real part of *ω*) with respect to the wave vector *k* determines the group velocity of the plasmon-polariton modes. The results are presented in Fig. 5. We observe that for the ionic system parameters listed above, the group velocity of the plasmon-polaritons reaches 100 m/s for the longitudinal mode, assuming that the initial post-synaptic AP or that from the axon hillock predominantly excites the longitudinal ion oscillations.

**Figure 5:**
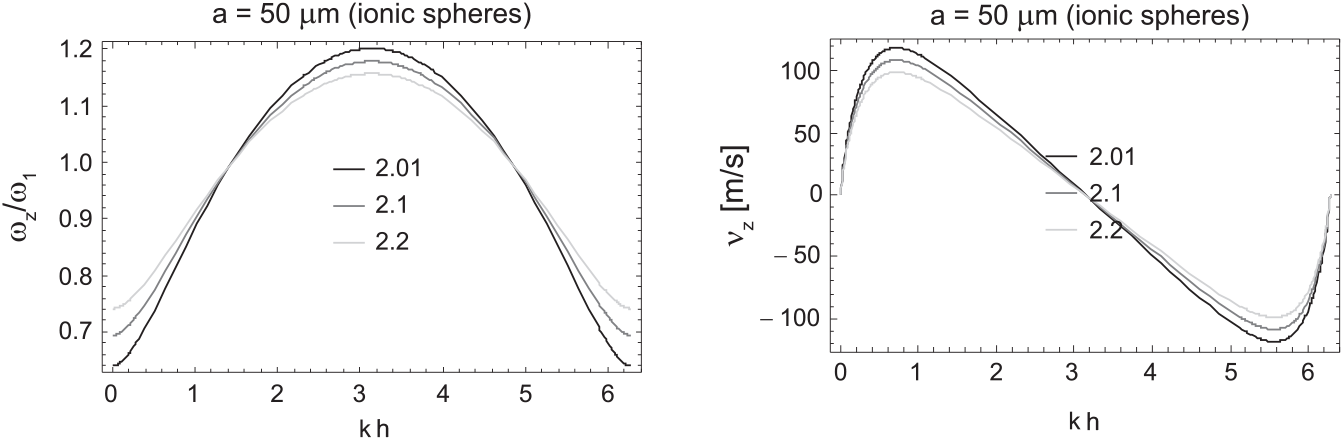
Solutions for the self-frequencies and group velocities of the longitudinal mode of a plasmon-polariton in the model ionic chain; *ω* is presented in units of *ω*_1_, here *ω*_1_ = 4 × 10^6^ 1/s, for a chain of spheres with radius *a* = 50 *μ*m and the Ranvier separation *d*_0_, *d*_0_/*a* = 2.01, 2.1, or 2.2, for an equivalent ion concentration in the inner cord of the axon of *n′* ~ 10 mM.

Note that similar plasmon-polariton kinetics occurs in finite chains of myelinated segments as in infinite chains, due to very fast convergence of the sums in Eq. (7) with denominators *m*^2^ and *m*^3^ (practically, a chain consisting of only 10 segments exhibits almost the same properties as an infinite chain).

In the model, the propagation of a plasmon-polariton through the axon chain and excited by an initial AP on the first node of Ranvier (after the synapse or, for the reverse signal direction, in the neuron cell hillock) sequentially triggers HH cycles at the consecutive Ranvier nodes with blocks of *Na*^+^ and *K*^+^ ion gates. The resulting firing of the AP traverses the axon with the velocity of ca. 100 m/s (or higher for thicker myelin sheath), consistent with the velocity actually observed in the myelinated axons. The plasmon-polariton ignition of the consecutive nodes of Ranvier releases the creation of the same AP pattern aided by the external energy supply at each node of Ranvier. Because of the nonlinearity of the HH mechanism (1–3), the signal initial growth saturates at a constant level, and the overall timing of each AP spike has a stable shape. The permanent supply of energy associated with creation of the AP spikes at sequentially firing nodes of Ranvier contributes also to the plasmon-polariton dynamics. This regeneration of the AP at each node of Ranvier assures that the dipole oscillation amplitude of plasmon-polariton is still beyond the activation threshold, despite Ohmic losses (actually rather small). The external energy supply (through the conventional ATP/ADP cell mechanism) assisting the renovation of the steady state of each node of Ranvier, residually compensates also the Ohmic thermal losses of the plasmon-polariton mode propagating along the axon and ensures undamped signal propagation over an unlimited range. Although the entire HH cycle of the AP spike on a single node of Ranvier requires several milliseconds (or even longer when one includes the time required to restore steady-state conditions, which, on the other hand, conveniently blocks the reversing of the signal propagation), subsequent nodes are ignited more rapidly, corresponding to the velocity of the plasmon-polariton wave packet triggering the ignition of the consecutive nodes of Ranvier, as illustrated in the Fig. 6. Thus, we deal with the firing of the full axon, where the ignition signal propagates with the group velocity of the plasmon-polariton wave packet.

**Figure 6:**
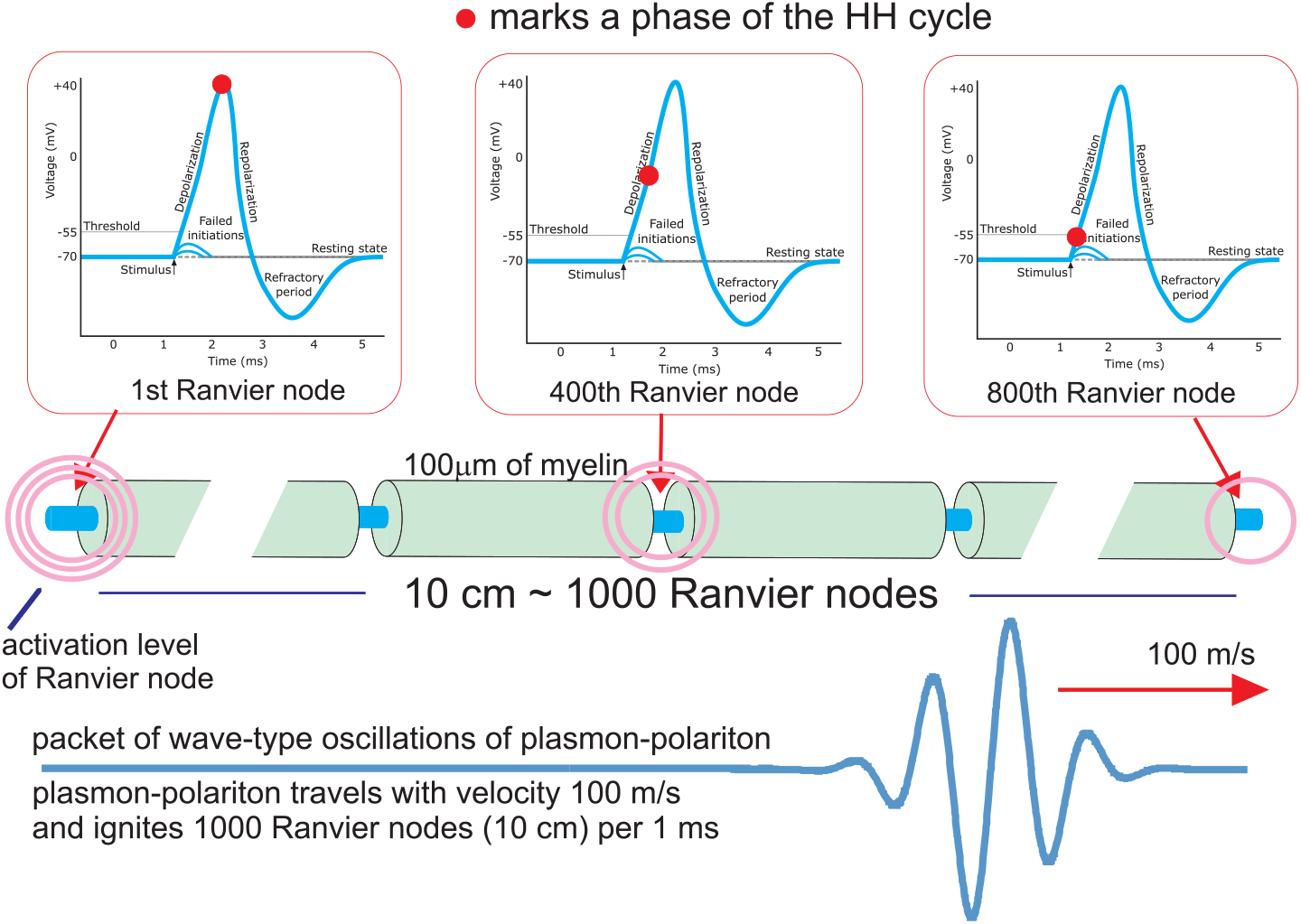
Schematic presentation of the firing of 10 cm long myelinated axon. The wave-packet of plasmon-polariton oscillations travels along the axon with the velocity of 100 m/s and in time of 1 ms ignites 1000 Ranvier nodes (as the myelinated segments have a length of 100 *μ*m each). The Hodgkin-Huxley (HH) cycle of the AP spike creation takes ca. 1 ms, thus consecutive Ranvier nodes are in various phases of HH cycle, as indicated by red points. Hence, the whole 10 cm long axon is in firing within the time period of 1 ms.

The direction of the velocity of the plasmon-polariton wave packet is adjusted to the semi-infinite geometry of the chain (in fact the chain is finite and is excited at one of its ends). The firing of the axon triggered by the plasmon-polariton traverses along the axon only in one direction because at nodes that had already ignited their Na^+^, K^+^ gates are inactivated (discharged) and require a relatively long time to restore the original steady status (the sodium/potassium block requires even one second time period and sufficient energy supply to bring Na^+^ and K^+^ ion concentrations to their normal values via cross-membrane active ion pumps against the ion-density gradient).

The firing of the AP can simultaneously move in two opposite directions if a certain central node of Ranvier of a passive axon is ignited, which has been observed. It is also observed the maintenance of the firing signal traversing despite small breaks in the axon cord or with even damaged few nodes of Ranvier, which agrees with the collective wave-type plasmon-polariton model of saltatory conduction in contrast to the lack of satisfactory explanations in the models based on the diffusive cable theory (4). The maintenance of the plasmon-polariton kinetics despite discontinuities in the axon cord agrees well with the discrete chain model of the cord corrugated by the periodic myelin sheath.

In Fig. 7 the group velocity of the AP traversing a firing myelinated axon (in the plasmon-polariton model) is plotted for various diameters of the axon internal cord, with the length of 100 *μ*m for each myelinated sector wrapped by Schwann (or oligodendrocyte) cells and Ranvier intervals of 0.5 *μ*m, 5 *μ*m, and 10 *μ*m width. The dependence of the group velocity on the width of the Ranvier interval is weak (i.e., negligible at the scale considered, which is consistent with the equivalence of the discrete model and the continuous system if one considers wave type plasmon-polariton propagation), but the increase in the velocity with increasing internal cord thickness is significant (the speed scales linearly with the axon diameter, i.e., is proportional to *d* = 2*r*), similarly to observed saltatory conduction in axons with increasing diameters.

**Figure 7:**
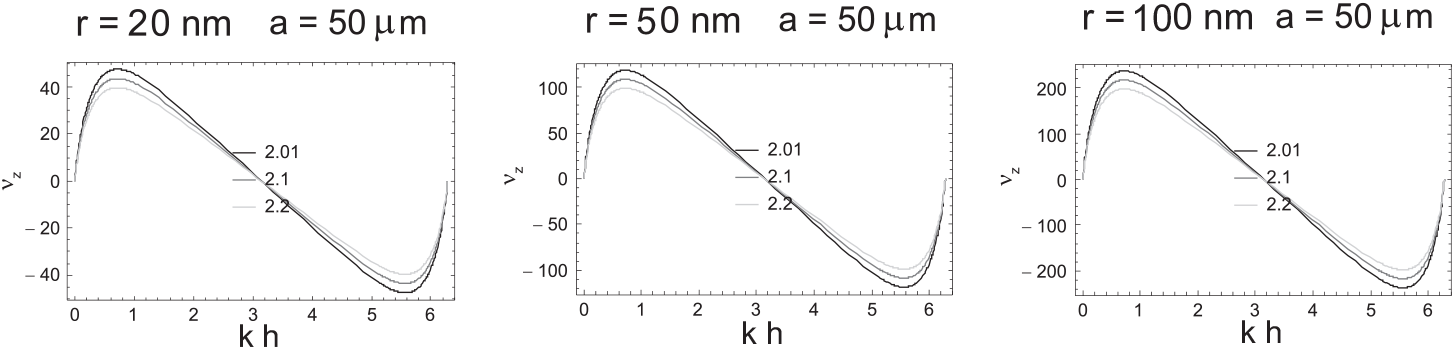
Comparison of the group velocities, in units of m/s, of the longitudinal plasmon-polariton mode with respect to the wave vector *k* ∈ [0, 2*π*/*h*) within the axon model for myelinated sectors with a length, *l* = 2*a* = 100 *μ*m and Ranvier node separation, *d*_0_ = 0.5 *μ*m, 5 *μ*m, and 10 *μ*m (represented by *d*_0_/*a* = 2.01, 2.1, and 2.2 in the figure, respectively) and for the axon cord radii of *r* = 20, 50, and 100 nm.

To comment on the appropriateness of the chain model for axons, let us note that even though the thin core of the axon is a continuous ion conducting fiber, the electromagnetic field can be closely pinned to the linear conductor surface similarly to the Goubau line (well known from microwave technology) (41, 42). For plasmon-polariton kinetics, the continuity of the conducting fiber is unimportant because the traversing of the wave packet of the plasma oscillations is not any current of charge, similarily to the Goubau microwave lines, which also have discontinuous segments. The Goubau lines maintain their transmittance via discrete disconnected elements. Synchronized oscillations of ion local concentration in chain segments when plasmon-polariton propagates do not give any net current of charge along the axon.

The fragments of the thin axon cord wrapped with thick myelin sheath with an ion concentration that is typical for neurons of *n′* ≃ 10 mM ≃ 6 × 10^24^ 1/m^3^ inside the cell can be equivalently modeled by a sphere with a diameter equal to the length of the cord fragment and the ion concentration of *n* ≃ 2 × 10^14^ 1/m^3^ (for an assumed axon cord diameter of 100 nm), conserving the number of ions in the segment participating in the dipole oscillations. Such a model is justified by the same structure of the dynamics equation as that for the dipole plasmon fluctuations in a chain of spheres and the modification of this equation for prolate spheroid or elongated cylindrical rod chains.

Because the dynamics equation is not affected by the anisotropy, the solutions of the equation for each polarization have the same form as for the spherical case. Thus, we can independently renormalize the equation for dipole oscillations for each polarization direction, introducing the resonance oscillation frequency for each direction *ω*_*α*1_ in a phenomenological manner (these frequencies can be estimated numerically, whereas for a sphere, 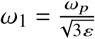; in general, the longer the semiaxis, the lower the related dipole oscillation frequency is).

The plasmon-polariton kinetics is very efficient and energetically economical signaling despite the rather poor ordinary electric conductivity of axons. Plasmon-polaritons do not radiate any energy. The energy supply to cover Ohmic losses is residually provided by the ATP/ADP mechanism at nodes of Ranvier, which energetically contributes to the restoration of the steady-state ion concentrations, via the active transport of ions across the membrane against the ionic concentration gradient, which requires an external energy supply. Moreover, the coincidence of the micrometer scale of the periodic structure of Schwann (or oligodendrocyte) cells in an axon (of approximately 100 *μ*m in length) with optimal requirements for the size of ionic chains additionally also supports the plasmon-polariton model.

It must be emphasized that plasmon-polaritons do not interact with external electromagnetic waves or, equivalently, with photons (even with adjusted frequency), which is a consequence of the large difference between the group velocity of plasmon-polaritons and the velocity of photons 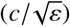. The resulting large discrepancy between the wavelengths of photons and plasmon-polaritons of the same energy prohibits mutual transformation of these two types of excitations because of the momentum-energy conservation constraints. Therefore, plasmon-polariton signaling by means of collective wave-type dipole plasmon oscillations along a chain, i.e., plasmon-polaritons, can be neither detected nor perturbed by external electromagnetic radiation. This also fits well with neuron signaling properties in the PNS and in the white myelinated matter in the CNS. The ionic surface plasmon frequencies are independent of the temperature. However, the temperature influences the mean velocity of ions, 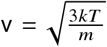, thereby enhancing the Ohmic losses with increasing temperature (cf. Eq. (3)), which in turn strengthens plasmon-polariton damping. Hence, at higher temperatures, higher external energy supplementation is required to maintain the same long-range propagation of plasmon-polaritons with a constant amplitude. This property is also consistent with experimental observations. The energy cost is the active ion transport through the membrane to recover the steady state of each node of Ranvier. Only a residual part of this energy is enough to eventually support plasmon-polariton amplitude at each node of Ranvier. Including this forcing field the plasmon-polariton can be considered in the regime of the steady solution of the forced and damped oscillator. The similar concerns plasmon-polariton dynamics in a metallic chain supplementary activated by a coupled external source (e.g., a coupled quantum dot system (43)). The speed of the plasmon-polariton signal transduction is temperature independent in contrary to cable theory discrete diffusion models (7, 12), the higher temperature causes, however, damping of plasmon-polariton.

The renovation of the AP at each Ranvier node assures the constant amplitude of the AP spike and, on the other hand, causes a convenient narrowing of the plasmon-polariton *k*-wave-packet, step-by-step supplementary re-excited at the consecutive Ranvier nodes. This narrowing shifts the packet toward the long-wave limit (small *k*), which follows from the Fourier picture of the Dirac delta (the larger number of the chain segments contributes to the excitation of the plasmon-polariton the narrower plasmon-polariton *k*-wave packet is formed). This favorable property allows for the precise ignition of the subsequent nodes of Ranvier by unlimited propagation of the relatively narrow plasmon-polariton wave packet precisely triggering consecutive nodes, which in turn feed the packet due to HH cycles.

## COMPARISON WITH THE CABLE MODEL

The cable theory is in fact the conventional model of a transmission line widely applied in the electronics and communication. The long line (like a coaxial line, particularly well corresponding to a bare neuron cell cord separated by a thin insulator membrane from the outer electrolyte of inter-cellular cytosol) is assumed to be a series of infinitely short segments of length *dx*, represented by the scheme as in Fig. 8. *R* is the longitudinal resistance of the inner line per its length unit, *L* is the distributed inductance of the inner wire per its length unit, *C* is the capacitance between two line components across the separating dielectric layer also per unit length of the line, *G* is the electrical conductivity across the barrier per unit length of the line.

**Figure 8:**
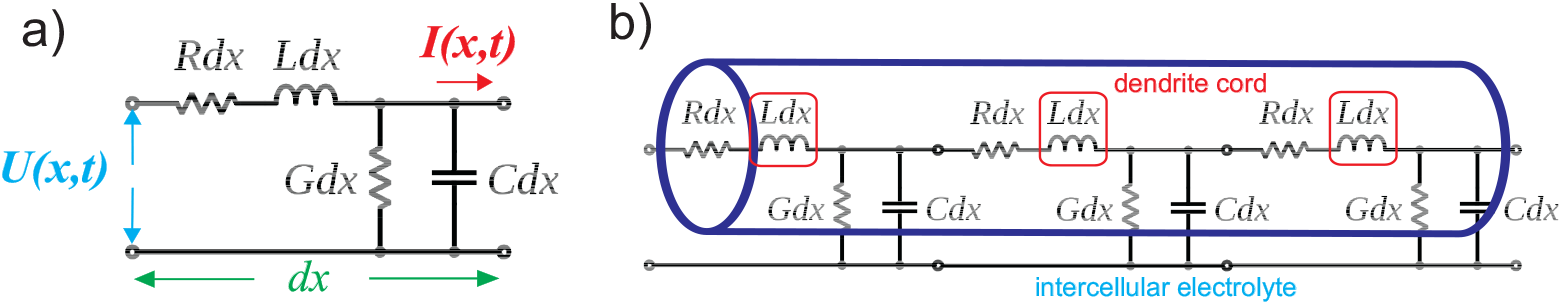
a) Elementary segment of the transmission line of length *dx*. *R* and L are resistance and inductance of the inner wire (the upper line in the figure) per length unit, *C* and *G* is the capacity and conductivity across the insulating barrier per unit of the length of the line. The electrical current flows only along the inner wire. b) Transmission line applied to a dendrite—the inner wire is the dendrite cord (with *L* = 0) and outer coaxial screen is the intercellular electrolyte separated by the dendrite (or unmyelinated axon) cell membrane.

Due to the Ohm law and the definition of the capacitance and the inductance one gets the relations between the voltage gated the line segment and the current along the inner wire,

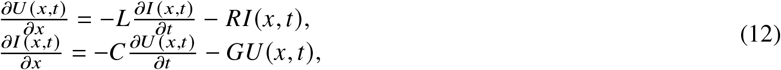

which are two coupled differential equations for the complex voltage *V* and current *I*. Taking the derivative 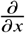 of both equations and substituting one into another, we can arrive in the second order differential equations,

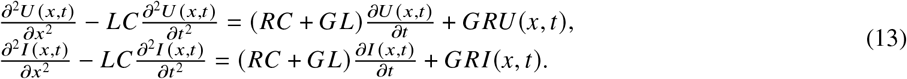

One can notice that both equations ate identical (they are conventionally called as telegrapher’s equations). For the lossless case, i.e., when *R* = *G* =0, Eqs (13) gain the form of the wave equations both for *U* and *I*, describing the ideal wave-type transmission along *x* direction with the velocity 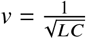 and without any losses. This corresponds to an ideal coaxial line. This is, however, not the case of a neuron, for which the assumption *L* = 0 is appropriate. Then Eqs. (13) attain the form of 1D diffusion equation,

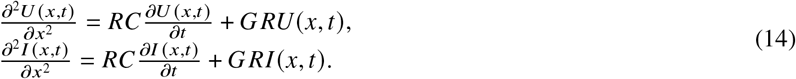

Defining the parameters 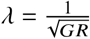 and 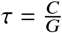, one can rewrite the above equation (for *U*) in a conventional form,

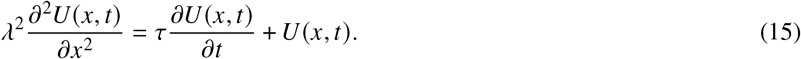

The parameter *λ* defines the spatial scale of the diffusion, whereas *τ* its time scale. The velocity of the diffusion of the signal, i.e., of the diffusive current along the dendrites (or unmyelinated axon) is assumed as 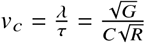. This velocity is the larger the smaller *C* and *R* and the larger *G* are. The range of the diffusion, ~ *λ*, lowers, however, with growing *G* (due to the shunt escape of the current). Larger *G* results in larger velocity but severely limits the range. The overall behavior of the *U*(*x*, *t*) (or *I*) diffusion defined by Eq. (15) is illustrated in Fig. 9, which presents the solution of Eq. (15) for initial condition in the form of periodic excitation in *x* = 0 point, *U*(0, *t*) = *U*_0_*cos*(*t*). One can notice that the cable theory (i.e., Eq. (15)) gives the non-wave-type propagation related to an ordinary current (and voltage signal) of diffusion type, thus on a relatively short distance with a quickly lowering amplitude. For the realistic values of *R*, *C*, *G* and d in axons, the estimation of *v_c_* gives 0.5 - 1 m/s. It is too low for explanation of the saltatory conduction in myelinated axons.

**Figure 9:**
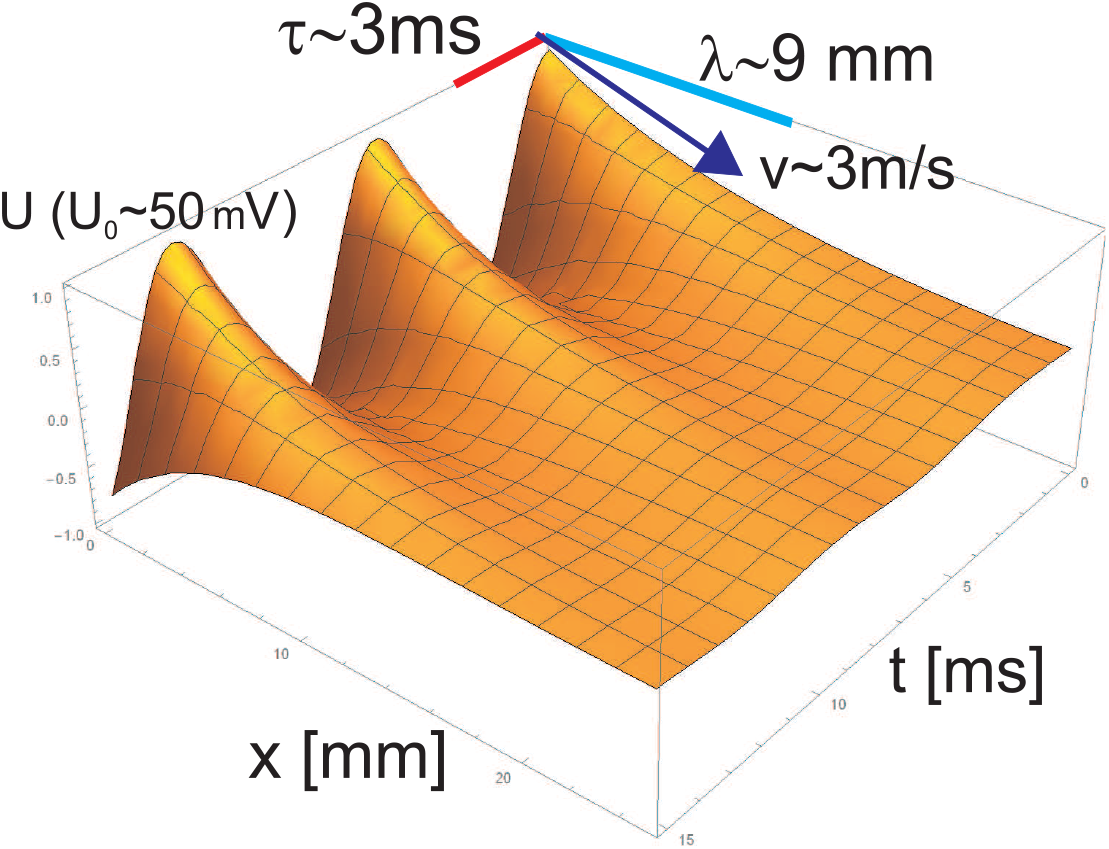
The solution of Eq. (15) for *λ* = 9 mm and *τ* = 3 ms and the ignition signal at *x* = 0 point, *U*(0, *t*) = 50*cos*(*t*) *mV*. The unit proportions in the figure are conserved.

The observed conduction velocity of a myelinated axon is by factor 100 – 200 greater than that of a geometrically similar unmyelinated axon. Myelinated neurons make up a large proportion of all neurons in the human body, more so in the CNS. Specifically, all PNS neurons with diameters greater than around 1 *μ*m and all CNS neurons with diameters greater than around 0.2 *μ*m are myelinated (9–11). In a study of myelinated axons in the human brain, most were found to have a diameter less than 1 *μ*m. The cable theory applied to myelinated axons gives at most 6 times larger velocity and to consider velocity ~ 40 m/s the axon diameter and internodal length are assumed extremely large (6) in contrary to observations (6, 9, 10). Data indicate that the *v_m_* is roughly proportional to fiber diameter, *d*, but this scale factor does vary. The complex factors impacting the *v_m_* in myelinated axons can be found in (9, 11) upon the cable theory variants, including, for instance, myelin thickness and capacitance (taking into account that the cable theory speed *v_c_* scales as 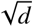). In the plasmon-polariton model we have got *v_m_* ~ *d*.

In discrete diffusion model (1) or in other modifications of the cable theory (2, 3) the HH cycle is included as the integrative element of local electricity in the closest adjacent myelinated segments. In the plasmon-polariton model the HH cycles at consecutive nodes of Ranvier are decoupled of the synchronic oscillations of ion density in myelinated segments. The HH cycles are triggered by the plasmon-polariton wave packet traveling along the axon with high speed. HH cycles play, however, a role of the control over plasmon-polariton via the resonance selection of particular modes which are supplemented in energy and thus traversing arbitrary long distances with the constant amplitude despite Ohmic losses at ion oscillations.

## COMMENTS AND CONCLUSIONS

We have provided a model for the saltatory conduction based on the kinetic properties of collective plasmon-polariton modes propagating along linear and periodically myelinated axons. Plasmon-polaritons were previously investigated and understood upon the very well developed nanoplasmonics of periodic metallic structures. The main properties of these collective excitations in metallic chains are,

- much lower velocity in comparison to the velocity of light, which gives much shorter wavelength of plasmon-polaritons than light wavelength with the same frequency,
- absence of radiative losses and long range almost undamped propagation.

These properties share also ion plasmon-polaritons in micro-chains of liquid electrolytes confined to a finite volume by appropriately formed membranes. Ionic plasmons and plasmon-polaritons have the frequency (energy) much lower than in metals and spatial micrometer size-scale instead of nanometer one in metals, both conveniently accommodated to living bio-cell organization because of higher mass of ions and their lower concentrations in cell cytosol in comparison to electrons in metals. Velocity of ionic plasmon-polariton can also be strongly reduced to the level just as for saltatory conduction in myelinated axons.

In a linear chain of ionic systems, the radiation losses of synchronized ion oscillation in chain segments in the form of plasmon-polariton are completely quenched to zero in exactly the same manner as in metallic chains. The high radiation energy losses expressed by the Lorentz friction at each chain element are perfectly compensated by the irradiation from the rest of the chain. Since the radiative losses are ideally balanced, only relatively low irreversible Ohmic energy dissipation remains. The collective plasmon-polaritons can thus propagate in the ionic linear periodic alignment with strongly reduced damping as in almost perfect wave-guide. If the energy is slightly supplemented to compensate the low Ohmic losses, the propagation of plasmon-polariton can reach arbitrarily long distances without any damping, which is observed in the saltatory conduction in myelinated axons. Moreover, the plasmon-polaritons in these axons cannot be neither detected nor perturbed by external e-m fields, due to a large incommensurability of plasmon-polariton and photon momenta at the same energy. Plasmon-polaritons may be thus carriers for triggering of the AP forming at consecutive nodes of Ranvier according to the HH cycle, in myelinated axons.

The advantages of the plasmon-polariton kinetics that the myelinated neuron signaling benefits from are following:

- Plasmon-polaritons cannot be perturbed nor detected by e-m waves (EEG/EMG signals are related to gray matter activity with ordinary conduction [according to the cable theory] which induces external e-m field signature, whereas myelinated white matter does not generate any external e-m signals).
- Immunity to e-m perturbation is convenient for reliable signaling in long links of myelinated neurons in the PNS (motorics and sensing cannot be disturbed by external e-m fields).
- All the e-m field of plasmon-polaritons is compressed along the tight tunnel around the axon cord.
- The myelin sheath must be sufficiently thick not for insulation but for the creation of the dielectric tunnel in the surrounding inter-cellular electrolyte to allow for plasmon-polariton formation with precisely tuned (by the meylin thickness) frequency and velocity of plasmon-polariton wave packet. Reducing of the myelin sheath thickness perturbs the synchronization of the plasmon-polariton frequency with the triggering time scale of Na^+^channels at Ranvier nodes, plasmom-polaritons start to be too quick for HH cycles and are damped because are not regenerated by HH cycles out of synchronization, the signal weakens and the group velocity of plasmon-polariton packet decreases if shifted to higher frequency (due to specific behavior of the group velocity of plasmon-polariton modes). This scenario takes place probably at Multiple Sclerosis or at other demyelination syndromes.
- Increasing temperature causes increase of Ohmic losses but does not reduce the plasmon-polariton group velocity. However, more energy is needed to compensate the increased Ohmic losses at higher temperatures, which might not be sufficiently supplemented by HH cycles.
- None radiative losses of Ohmic type occur only for plasmon-polaritons; if plasmon-polaritons are residually aided by AP spike formation (renovation upon the HH cycles) at consecutive nodes of Ranvier, an arbitrary long range plasmon-polariton kinetics is possible with constant dipole signal amplitude enough to trigger all HH cycles in series.
- The plasmon-polariton group velocity increases proportionally to the diameter of the axon cord.
- The plasmon-polariton group velocity for real neuron electrolytes and axon size parameters fits to the observed velocity of the saltatory conduction.
- Wave type plasmon-polariton kinetics explains the maintenance of signal transduction despite small breaks and gaps in the neuron cord or a damage of some nodes of Ranvier (these observations were impossible to be explained within the cable theory or by the compression-soliton model).
- One-direction kinetics of plasmon-polariton occurs if started from one end of an axon, but two-direction kinetics occurs if plasmon-polariton is ignited from a certain central point of a passive axon in agreement with observations.

Taking into account the properties of plasmon-polariton conduction listed above one might thus propose a nonlocal-topological braid-group-type approach to information processing in the brain considering the specific and different electricity of the gray and white matter of neurons. The web of neuron filaments in gray matter would serve to encode the information in the entanglement of filaments selected by activation of certain synapses, because the electricity of dendrites and of unmyelinated axons (in the gray matter) is of an electrical current type providing an e-m signature when an AC signal of brain waves is transduced through the whole neuron network. The method of e-m resonance identification of information stored in braided segments of the neuron web can be thus suggested. This method favors the gray matter with ordinary ion currents of the diffusive type (as in the cable model) but excludes the myelinated white matter with wave-type plasmon-polariton signaling without any e-m signature. The saltatory conduction in myelinated axons in the white matter is not active with respect to the e-m resonance, which property is typical for plasmon-polaritons. This wave type propagation of the AP is nonlocal in contrast to diffusion model (upon the cable theory) and conveniently fulfills a communication role in the PNS and CNS. The plasmon-polariton model explains the efficient, very fast and long range saltatory conduction in myelinated axons in the PNS and in the white matter of the brain and spinal cord, where the velocity and the energy-efficiency of communication is of the highest significance. The close correspondence with observations supports the propriety of the new model proposed for saltatory conduction of the AP, which, on the other hand, complementary fits with the different role and functioning of the gray and white matters.

## ACKNOWLEDGMENTS

Supported by NCN project P.2018/31/B/ST3/03764.

